# From APOE Genetics to AI-Designed Drug Candidates: An Integrated Pipeline for Oral, Brain-Penetrant ACAT1 Inhibitors in Alzheimer’s Disease

**DOI:** 10.64898/2026.05.21.727002

**Authors:** Sharad Agarwal, Rory Popert, Paul Agapow, Crystal Ruff, Shikta Gupta

## Abstract

For more than 55 million people living with dementia worldwide, no oral disease-modifying treatment is currently available. Alzheimer’s disease (AD) remains one of the most urgent unmet needs in neurology, with recent genetic evidence estimating that 72–93% of AD burden is attributable to common APOE allelic variation. Mechanistically, the APOE ε4 isoform impairs cholesterol transport in the brain, promoting cholesteryl ester accumulation in microglia; these lipid-laden cells lose phagocytic capacity for amyloid-β clearance and autophagy-mediated degradation of phosphorylated tau, linking a single upstream metabolic disruption to both hallmark pathologies of AD. ACAT1/SOAT1, the brain’s predominant cholesterol-esterifying enzyme, is an attractive therapeutic target: its inhibition reduces cholesteryl ester formation, restoring microglial Aβ clearance, reducing Aβ production, and promoting tau degradation via autophagy, supporting a multimodal mechanism not addressed by currently approved therapies. However, prior ACAT inhibitor programs were limited by isoform non-selectivity, excessive lipophilicity, and lack of CNS optimization. Addressing these constraints, we present the CuraGenAI Drug Discovery Platform for the design of oral, brain-penetrant ACAT1-targeted small-molecule candidates. The pipeline integrates scaffold-constrained generative chemistry with multi-objective ADMET optimization across more than 30 criteria. From approximately 6 million generated molecules, multi-stage filtering yielded approximately 7,300 CNS-optimized candidates with predicted oral brain-penetrant profiles distinct from prior ACAT clinical compounds. Three nominated lead candidates showed predicted BBB probabilities >0.93, favorable CNS drug-like physicochemical profiles, and stable ACAT1 binding poses, and are prioritized for synthesis and in vitro validation.

## 1. Introduction

### 1.1 The Alzheimer’s Disease Treatment Gap

Alzheimer’s disease remains the leading cause of dementia worldwide; collectively, dementia affects more than 55 million people with a projected economic burden exceeding $1 trillion annually [1]. The drug development landscape is defined by an extraordinary failure rate: approximately 98% of AD clinical trials initiated since 2003 have failed. Currently approved therapies fall into two primary categories: 1) symptomatic treatments that provide modest and temporary cognitive benefit, and 2) the recently introduced anti-amyloid monoclonal antibodies lecanemab [2] and donanemab [3], which represent the first disease-modifying agents.

The anti-amyloid antibodies target and clear amyloid plaques from the brain, slowing cognitive decline by 27–35% [2, 3], leaving substantial room for complementary oral disease-modifying approaches. Despite this novel mechanism, these biologics also carry significant risk of amyloid-related imaging abnormalities (ARIA), occurring in 12.6–24% of treated patients and disproportionately affecting APOE ε4 carriers, the very population at highest AD risk. The treatment burden is also substantial: intravenous infusion every two weeks, mandatory MRI monitoring, and annual costs of $26,000–$32,000 per patient. These constraints create a clear unmet need for oral, disease-modifying therapies that target upstream AD pathology through mechanistically distinct pathways not expected to carry antibody-associated ARIA risk.

### 1.2 APOE and the Cholesterol–Alzheimer’s Connection

The apolipoprotein E (APOE) gene has been recognized as the strongest genetic risk factor for late-onset AD since 1993. A landmark study published in January 2026 by Williams et al. [4] estimated that 72–93% of AD cases would not occur without the combined influence of the APOE ε3 and ε4 alleles. This finding redefines the therapeutic opportunity: rather than targeting downstream amyloid aggregates, interventions that target the upstream cholesterol dysregulation driven by APOE may offer disease-modifying potential.

The mechanistic link between APOE genotype and AD pathology converges on cholesterol homeostasis in the brain [5, 6]. APOE is the primary cholesterol transporter in the CNS; the ε4 isoform is structurally less efficient at mediating cholesterol efflux and lipoprotein receptor-mediated clearance, promoting cholesteryl ester accumulation in neurons and glia [5, 6]. Cholesteryl ester-laden microglia lose phagocytic capacity for amyloid-β (Aβ) clearance and autophagy-mediated degradation of phosphorylated tau [14, 15], and neuronal cholesterol excess promotes amyloidogenic processing of APP (amyloid precursor protein), linking a single upstream metabolic disruption to both hallmark pathologies of AD.

Recent single-cell transcriptomic studies have further clarified the link between cholesterol dysregulation and microglial dysfunction. Cholesteryl ester accumulation drives formation of lipid droplets in microglia, promoting a transition to inflammatory, debris-loaded phenotypes with impaired phagocytic clearance capacity [7]. These lipid-droplet-accumulating microglia (LDAM) are enriched in APOE ε4 carriers and in brain regions with high amyloid burden, establishing lipid handling, particularly the cholesterol esterification pathway catalyzed by ACAT1, as a central node in the microglial immune response to AD pathology rather than a peripheral metabolic bystander.

### 1.3 ACAT1: The Mechanistic Convergence Node

Acyl-CoA:cholesterol acyltransferase 1 (ACAT1/SOAT1) is an endoplasmic reticulum-resident integral membrane enzyme that catalyzes the esterification of free cholesterol to cholesteryl esters. ACAT1 is the predominant cholesterol-esterifying enzyme in the brain. It belongs to the membrane-bound O-acyltransferase (MBOAT) superfamily [8], sharing a conserved catalytic histidine residue with ten other human enzymes.

Building on foundational ACAT1/SOAT1 biology from the Chang laboratory and others [9, 10], ACAT1 is a historically proposed but clinically underexplored AD target whose rationale has strengthened through recent evidence from APOE genetics, lipid-droplet microglia biology, and TREM2/sTREM2/LRP1 signaling [4, 7, 12]. Because ACAT1 controls the balance between free cholesterol and cholesteryl ester storage in the brain, its inhibition reverses the cholesteryl ester accumulation described above, with preclinical evidence across three experimentally tractable mechanisms of action:

#### Aβ reduction

Bryleva et al. [11] demonstrated that ACAT1 gene ablation in AD-model mice reduced APP levels by over 60%, nearly eliminated amyloid plaque burden, and restored cognitive function.

#### Microglial phagocytic restoration via sTREM2/LRP1

Hovde, Tanzi and colleagues [12] demonstrated that ACAT1 inhibition promotes shedding of soluble TREM2 through α-secretase activation in microglia, facilitating LRP1-dependent microglial uptake of Aβ.

#### Tau reduction via autophagy

Huynh et al. [13] showed that ACAT1/SOAT1 inhibition in aging APOE ε4 mice reduced brain neuroinflammation, and Shibuya et al. [14] demonstrated that ACAT1 inhibition in microglia stimulates autophagy-mediated lysosomal proteolysis with evidence of enhanced clearance of tau aggregates. Consistent with this tau-linked mechanism, human iPSC-derived neuron studies have identified cholesteryl ester accumulation as an upstream regulator of phosphorylated tau through proteasome-dependent pathways [15].

The therapeutic rationale is reinforced by orthogonal interventions: pharmacological ACAT inhibition reduced amyloid plaques by 88–99% in APP transgenic mice [16], AAV-mediated Acat1 knockdown reduced brain Aβ within two months even in symptomatic 3xTg-AD mice, suggesting efficacy beyond prevention into established disease [17], and selective inhibition by K-604 increased microglial Aβ uptake and lysosomal volume at partial ACAT1 inhibition [14]. Together, these genetic, pharmacological, and gene-silencing data support a multimodal hypothesis in which amyloid production, microglial clearance, and autophagy-mediated tau clearance pathways may be modulated through a single target not addressed by currently approved AD therapies. Because ACAT1 is expressed in neurons, astrocytes, and microglia, an oral small-molecule inhibitor would engage the pathway across multiple brain cell types, reducing amyloidogenic APP processing in neurons [11] while restoring phagocytic clearance in microglia [12], rather than requiring cell-type-specific targeting. Each axis is linked to a quantifiable biomarker: sTREM2 shedding (ELISA [12]), Aβ phagocytosis rate (fluorescent uptake assay [12]), and cholesteryl ester/free cholesterol ratio (lipidomics), enabling pharmacodynamic monitoring from preclinical assays through clinical trials.

### 1.4 Lessons from Prior ACAT Inhibitor Programs

No ACAT1-selective inhibitor has ever entered clinical trials for AD. Prior clinical and preclinical ACAT inhibitor programs — avasimibe from Pfizer, eflucimibe from Pierre Fabre, and nevanimibe from Millendo — were ultimately discontinued [18] due to isoform non-selectivity, excessive lipophilicity driving peripheral tissue toxicity (LogP >6, adrenal foamy cell accumulation), and lack of CNS optimization. The sole partially selective compound, K-604 (Kowa), was developed for atherosclerosis, not AD. Prior programs did not incorporate off-target counter-screening, blood–brain barrier penetration constraints, or safety profiling for chronic oral dosing in elderly patients.

### 1.5 The Present Opportunity

Three developments create a unique window for revisiting ACAT1/SOAT1 as an Alzheimer’s disease target: (1) APOE genetic evidence implicating cholesterol metabolism as a central axis in AD pathology [4]; (2) the Hovde/Tanzi discovery that ACAT1 inhibition promotes microglial Aβ clearance through a novel sTREM2/LRP1-dependent pathway, strengthening the rationale for ACAT1-targeted multimodal disease-modification and providing a measurable pharmacodynamic axis [12]; and (3) the availability of experimentally determined ACAT1/ACAT2 cryo-EM structures and expanded historical ACAT inhibitor SAR, enabling structure-guided design of CNS-optimized candidates. We present an end-to-end computational pipeline, developed within the CuraGenAI Drug Discovery Platform, for the design of oral, brain-penetrant ACAT1-targeted small-molecule candidates for AD.

## 2. Methods

### 2.1 Computational Pipeline Architecture

The CuraGenAI generative engine employs a graph-based encoder–decoder transformer [19, 20] architecture for scaffold-constrained de novo drug design, in which the encoder receives molecular fragments as input scaffolds and the decoder iteratively predicts atom types and bond connectivities to generate complete structures. Molecules are represented as graph-encoded matrices in which rows encode atom types, bond types, and connectivity information, with positional encoding derived from the molecular adjacency matrix to capture graph topology within the transformer attention mechanism [19]. This graph-based representation was selected over SMILES-based alternatives because it enforces scaffold constraints during generation (preserving pharmacophoric fragments intact) and avoids syntactic validity issues inherent in string-based approaches.

The generative model was pre-trained on approximately 1.6 million drug-like molecules from a curated standardized bioactivity dataset [23] to learn the general grammar of drug-like chemical space. The pre-trained model was then fine-tuned on the 562 ACAT1-active compounds described in Section 2.3. During the subsequent reinforcement learning (RL) phase, the agent’s policy was updated using a multi-objective reward signal integrating four concurrent objectives: positive structure-based target engagement evaluated by AutoDock-GPU against ACAT1 (Protein Data Bank (PDB) 6VUM), an inverse docking penalty against ACAT2 (PDB 7N6R) for selectivity-by-design, pharmacokinetic and safety fitness computed by the CuraGenAI ADMET Profiling Module (Section 2.7), and synthetic accessibility [24].

The CuraGenAI scoring architecture separates training and generation modes. During training, molecules violating pharmacokinetic or safety constraints receive soft score penalties, enabling broader exploration of viable CNS drug space. During generation, the same constraints are enforced as hard binary filters: molecules failing any critical endpoint are excluded. This two-phase design prevents premature convergence to a narrow chemical series while ensuring stringent quality in the final output.

The pipeline was further customized for the ACAT1/Alzheimer’s application through five domain-specific additions: CNS-optimized ADMET scoring (Section 2.7), ICH M7-aligned structural alert filtering (Section 2.8), exhaustive stereoisomer enumeration (Section 2.9), MBOAT family counter-screening (Section 2.10), and dual-source fragment curation combining data-driven and expert-curated seeds (Section 2.5). All computations were performed on NVIDIA GPU hardware. Complete protocols are provided in Supplementary Methods.

### 2.2 Target Structure Selection

The primary docking target was PDB 6VUM, the 3.5 Å cryo-EM structure of human tetrameric ACAT1 in complex with nevanimibe [25], selected because the co-crystallized ligand defines a pharmacologically relevant binding pocket within the TM4–TM9 catalytic chamber containing His460. Receptor preparation used Reduce2 and Meeko. AutoDock-GPU was used for RL-stage and generation-scale scoring; final lead candidates were independently rescored using AutoDock Vina consensus docking (Section 3.5). Top-ranked candidates were validated by re-docking against PDB 6L47 (CI-976-bound ACAT1, 3.5 Å) [26] to confirm binding mode consistency.

For ACAT2 counter-screening, PDB 7N6R (nevanimibe-bound) served as the primary target and PDB 7N6Q as the validation structure. Docking scores were interpreted as relative binding energy estimates for compound ranking, not as absolute binding affinity predictions. Redocking nevanimibe into 6VUM reproduced the cryo-EM binding pose with an RMSD of 2.6 Å, within the expected range for 3.5 Å resolution cryo-EM data. Full protocol details are provided in Supplementary Methods.

### 2.3 Compound Dataset Assembly

ACAT1-active compounds were assembled from pharmacophore-based virtual screening of PubChem using Pharmit [27], curated bioactivity databases [23], and literature extraction of known ACAT1 inhibitors. The assembled library was scored by AutoDock-GPU against PDB 6VUM, retaining 562 compounds with confirmed docking engagement (scores from −13.66 to −8.33 kcal/mol, mean: −10.41 kcal/mol). The 562-compound dataset reflects the limited public bioactivity data available for ACAT1, an underexplored target compared to kinases or GPCRs with thousands of annotated compounds, and the pipeline accommodates this by deriving the primary RL reward from physics-based docking rather than from quantitative structure–activity relationship (QSAR) models that would require larger training sets (Section 4.2).

### 2.4 Murcko Scaffold Decomposition and Chemotype Classification

All 562 compounds were subjected to Murcko scaffold decomposition (RDKit), yielding 395 unique scaffolds and 322 generic frameworks. Scaffolds were classified into chemotype families by SMARTS matching against twelve reference pharmacophores from the ACAT1 inhibitor literature and assigned to three categories:

**• LEVERAGE** (primary generative seeds): Chemotypes with demonstrated ACAT1 activity or validated structural biology, including the K-604 benzimidazole-piperazine core [28], the CI-976 trimethoxyphenyl amide (co-crystallized in PDB 6L47), and CNS-privileged linker scaffolds.
**• AVOID** (excluded from fragment library): Chemotypes carrying liabilities intrinsic to the core scaffold: acyl sulfonamides (avasimibe class; dual ACAT1/ACAT2, CYP3A4 inducer, Phase III failed), disubstituted ureas (nevanimibe class; adrenal toxicity from lipophilicity), and pyranose scaffolds (pyripyropene derivatives; ACAT2-selective).
**• EXPLORE** (novel scaffolds for creative generation): All remaining compounds, representing the majority of the dataset and confirming that most validated ACAT1 binders occupy structurally unexplored space.

### 2.5 Dual-Source Fragment Curation

The fragment library was assembled through a dual-source strategy balancing exploitation of known ACAT1 pharmacophores with exploration of novel chemical space.

#### Source A (Data-Driven)

Murcko scaffolds from the highest-scoring compounds in each non-AVOID chemotype family (Section 2.4) were filtered for generative suitability: MW <300 Da, topological polar surface area (TPSA) <90 Å², maximum three per family. This yielded six data-driven fragments spanning four chemotype families.

#### Source B (Expert-Curated)

Twenty-one fragments were designed through iterative expert curation informed by structural analysis of the ACAT1 binding site (PDB 6VUM, 6L47), published selectivity SAR, and CNS-privileged scaffold literature. Design was guided by three criteria: (i) compatibility with the His460/Asn421 interaction geometry in the TM4–TM9 cavity; (ii) ACAT1/ACAT2 selectivity determinants, particularly the TM7/TM8 subpocket divergence; and (iii) CNS-privileged scaffolds with validated blood-brain barrier (BBB) permeability. Strategies included bioisosteric replacement to reduce HBD count (e.g., benzothiazole for benzimidazole), soft-drug lactone incorporation, and minimal pharmacophore deconstruction to retain pocket complementarity while eliminating lipophilicity-driven liabilities. Fragments comprised 7 LEVERAGE and 14 EXPLORE scaffolds.

The merged library (27 unique fragments: 9 LEVERAGE, 18 EXPLORE) was confirmed structurally distinct by pairwise Tanimoto similarity (Morgan fingerprints, radius 2, 2048 bits). Details are provided in Supplementary Table S2.

### 2.6 Generative Chemistry and Multi-Objective Optimization

Model training follows a three-phase transfer learning strategy implementing the architecture and reward described in Section 2.1. Phase 1 (pre-training): the model was trained on the approximately 1.6 million-compound bioactivity dataset [23] for 1,000 epochs with batch size 512 using the teacher forcing paradigm, learning the general grammar of valid molecular graphs. Phase 2 (fine-tuning): the pre-trained weights were adapted to ACAT1-relevant chemical space by 20 epochs of continued training on the 562 ACAT1-active compounds (Section 2.3) at a reduced batch size of 16 to bias toward ACAT1-relevant patterns while preventing catastrophic forgetting of general molecular grammar. Phase 3 (reinforcement learning): the fine-tuned model was further optimized against the multi-objective reward over 100 epochs, structured as a two-phase curriculum: an initial 14-epoch warm-up optimizing ACAT1 binding, ADMET, and synthetic accessibility, followed by 86 epochs with the ACAT2 counter-screening penalty active. With a sample size of 2,048 molecules per epoch, approximately 205,000 molecules were evaluated and scored during RL training.

### 2.7 ADMET and Safety Scoring

Each generated molecule was evaluated by the CuraGenAI ADMET Profiling Module across over 30 predicted pharmacokinetic, safety, and physicochemical endpoints organized for oral CNS drug candidates targeting chronic Alzheimer’s therapy (Supplementary Table S1).

The module integrates machine learning ensemble predictions [21, 22] with orthogonal physicochemical descriptors and structural alert screening in a consensus filtering strategy. Machine learning ADMET models are sensitive to training data composition and applicability domain, particularly for novel chemotypes. To mitigate single-model dependency, probabilistic predictions are complemented by RDKit-derived physicochemical descriptors (MW, LogP, TPSA, HBD, Fsp3, quantitative estimate of drug-likeness (QED) [29], rotatable bonds), RDKit FilterCatalog structural alerts (PAINS [30], Brenk [31]), and program-specific toxicophore SMARTS patterns (Section 2.8). This multi-source approach ensures that no single predictive component serves as the sole basis for any safety-critical decision, and that compounds advancing through the pipeline satisfy both statistical and rule-based (ICH M7 [32]) criteria simultaneously.

The composite ADMET fitness score was computed as a weighted combination of normalized endpoint desirability terms. Safety-critical endpoints with direct precedent in ACAT1 inhibitor clinical failures received the highest weights: BBB penetrance, hERG cardiac safety, drug-induced liver injury (DILI) risk, and CYP3A4 inhibition (addressing the avasimibe CYP induction failure). Multiplicative penalty functions were applied for molecules approaching hard-fail thresholds on critical endpoints, including low BBB penetrance, elevated P-glycoprotein efflux, CYP3A4 substrate liability, and unfavorable combinations of high lipophilicity with low hepatic clearance, specifically addressing the nevanimibe adrenal toxicity pattern (Section 1.4). Low Fsp3 was also penalized, as flat aromatic molecules are associated with promiscuous binding and poor solubility [33].

In generation mode, a binary hard-fail gate eliminated molecules exceeding absolute thresholds on approximately 35 critical endpoints, set based on published CNS drug space boundaries [34] and the clinical failure analysis in Section 1.4. Scoring weights, thresholds, and categories are summarized in Supplementary Table S1.

### 2.8 Structural Alert Filtering

A three-tier system was applied. Tier 1 (chemical instability) eliminated molecules containing O–F bonds, azides, or peroxides. Tier 2 (reactive and high-liability functionality) eliminated Michael acceptors, aldehydes, acyl halides, epoxides, alkyl halides, α-halo carbonyls, terminal alkynes, quinones, N-nitroso compounds, and hydrazines (ICH M7 cohort-of-concern structures [32]). Tier 3 (medicinal chemistry alerts) eliminated anilines and nitroaromatics. SMARTS-based pattern matching was used for all tiers to ensure deterministic flagging of structures that probabilistic ADMET predictors may not reliably identify.

### 2.9 Stereoisomer Enumeration

All surviving molecules underwent exhaustive stereoisomer enumeration, with each stereoisomer carried forward as an independently scored entity. Parent molecules without explicit stereo specification were removed, since racemic synthesis would yield mixtures with binding profiles potentially divergent from docking predictions. Only stereochemically defined entities were retained, ensuring all candidates are suitable for enantioselective synthesis.

### 2.10 MBOAT Family Counter-Screening Panel

ACAT1 belongs to the membrane-bound O-acyltransferase (MBOAT) family, whose members share a conserved catalytic histidine and transmembrane architecture [35, 36]. Four family members were selected for counter-screening based on the clinical consequences of their inadvertent inhibition in chronic oral Alzheimer’s therapy:

- **ACAT2** (PDB 7N6R, 7N6Q) [37]: inhibition depletes hepatocyte cholesteryl ester pools required for lipoprotein assembly: the mechanism that elevated LDL and contributed to avasimibe’s Phase III failure.
- **DGAT1** (PDB 8ESM, 8ETM) [38]: catalyzes enterocyte triglyceride synthesis; clinical DGAT1 inhibitors caused dose-limiting gastrointestinal (GI) intolerance, unacceptable for chronic dosing in elderly patients.
- **PORCN** (PDB 9OO6, 9OO7) [39]: palmitoylates all 19 human Wnt ligands; off-target inhibition blocks Wnt signaling required for adult hippocampal neurogenesis, which could be counterproductive in a chronic AD setting.
- **HHAT** (PDB 7Q6Z, 7Q1U) [40]: palmitoylates Hedgehog ligands required for subventricular zone neural stem cell maintenance; off-target inhibition risks impaired neurogenesis alongside bone loss and muscle wasting.

RL feedback operates on ACAT1 (6VUM) and ACAT2 (7N6R) only; remaining MBOAT off-targets are evaluated post-generation. Structure-based scoring uses AutoDock-GPU against cryo-EM structures.

### 2.11 Orthogonal Safety Confirmation of Lead Candidates

Lead candidates passing in-pipeline ADMET scoring, structural alert filtering, and MBOAT counter-screening underwent a final orthogonal safety confirmation. Regulatory-oriented safety predictions from VEGA (consensus mutagenicity, carcinogenicity, developmental toxicity) [43] and OPERA (Tox21-aligned ADME and toxicity endpoints) [44] were combined with CuraGenAI ADMET Module predictions (Section 2.7) and purpose-built Chemprop message-passing neural network (MPNN) ensemble classifiers for high-impact clinical attrition endpoints: hERG, DILI, BSEP inhibition (cholestatic DILI mechanism), and OATP1B1 inhibition (FDA-flagged DDI transporter relevant to polypharmacy in elderly AD patients). Candidates flagged consistently across all sources on any safety-critical endpoint were eliminated. Discordant calls triggered expert review to distinguish genuine liabilities from model noise. This orthogonal confirmation strategy explicitly targets the cardiac, hepatic, and drug–drug interaction liability classes that dominate Phase 1–3 clinical attrition.

## 3. Results

### 3.1 Dataset Characteristics and Chemotype Landscape

The reference dataset of 562 ACAT1-active compounds exhibited a significant constraint for CNS therapeutic development: only 6% (34/562) satisfied a composite CNS drug-likeness filter (MW <450, LogP <4, TPSA <90, HBD ≤2), with LogP the most restrictive criterion (only 114/562, 20%, below LogP 4). Dataset-wide properties confirmed pronounced lipophilicity bias (mean MW 464 Da, LogP 5.3, TPSA 81.0 Å²), reflecting the historical focus on peripheral cardiovascular targets rather than a fundamental incompatibility of the ACAT1 pharmacophore with CNS drug space. Chemotype classification (Section 2.4) distributed compounds into LEVERAGE (34, 6.0%), EXPLORE (460, 81.9%), and AVOID (68, 12.1%). Strikingly, 78% of the dataset (441/562) did not match any historically characterized pharmacophore family, indicating that the majority of validated ACAT1 chemical matter remains structurally unexplored.

### 3.2 Dual-Source Fragment Library

Source A (data-driven) yielded six fragments from four non-AVOID chemotype families satisfying MW <300 Da and TPSA <90 Å². Source B (expert curation) contributed 21 fragments: 7 LEVERAGE scaffolds with structural modifications for CNS properties, and 14 EXPLORE scaffolds from CNS-privileged chemical classes. The merged library comprised 27 unique fragments (9 LEVERAGE and 18 EXPLORE), occupying physicochemical space significantly different from the reference dataset: mean MW 199 Da (vs 464 Da), mean LogP 2.2 (vs 5.3), and mean TPSA 40.9 Å² (vs 81.0 Å²), providing CNS-compatible starting points while leaving approximately 150–250 Da of molecular weight budget for AI-driven elaboration toward final compounds of 350–450 Da. Fragment classification, scaffold class, and physicochemical property ranges are provided in Supplementary Table S2.

### 3.3 Generative Chemistry Output

The generative engine produced approximately 6 million molecules across iterative generation cycles. After sequential application of ADMET hard-fail filters (approximately 1% pass rate), exhaustive stereoisomer enumeration, and three-tier structural alert filtering, approximately 224,000 stereochemically defined candidates advanced to structure-based scoring. Of these, approximately 7,300 passed ACAT1 engagement and ACAT2 counter-screening gates, advancing to orthogonal safety confirmation.

### 3.4 ADMET Filtering Cascade: Enrichment and Attrition

The multi-stage filtering architecture eliminated compounds through three sequential stages, each targeting distinct liability classes (Figure 3).

**Figure 1.**
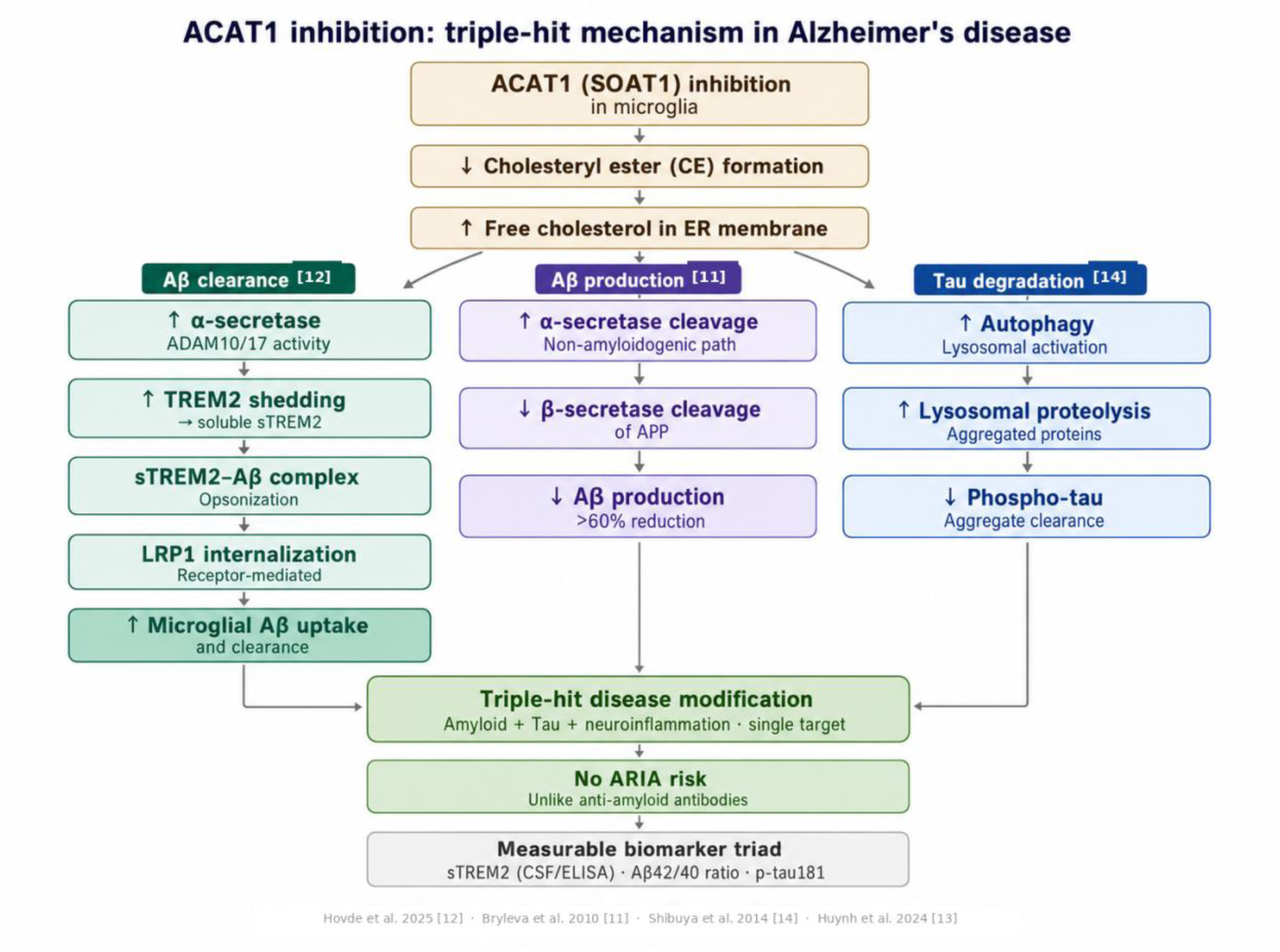
ACAT1 inhibition drives a triple-hit mechanism via three parallel pathways: microglial Aβ clearance (sTREM2/LRP1) [12], Aβ production reduction (>60%) [11], and phospho-tau degradation (autophagy) [14]. All converge on a single-target disease-modification hypothesis not expected to carry antibody-associated ARIA risk.

**Figure 2.**
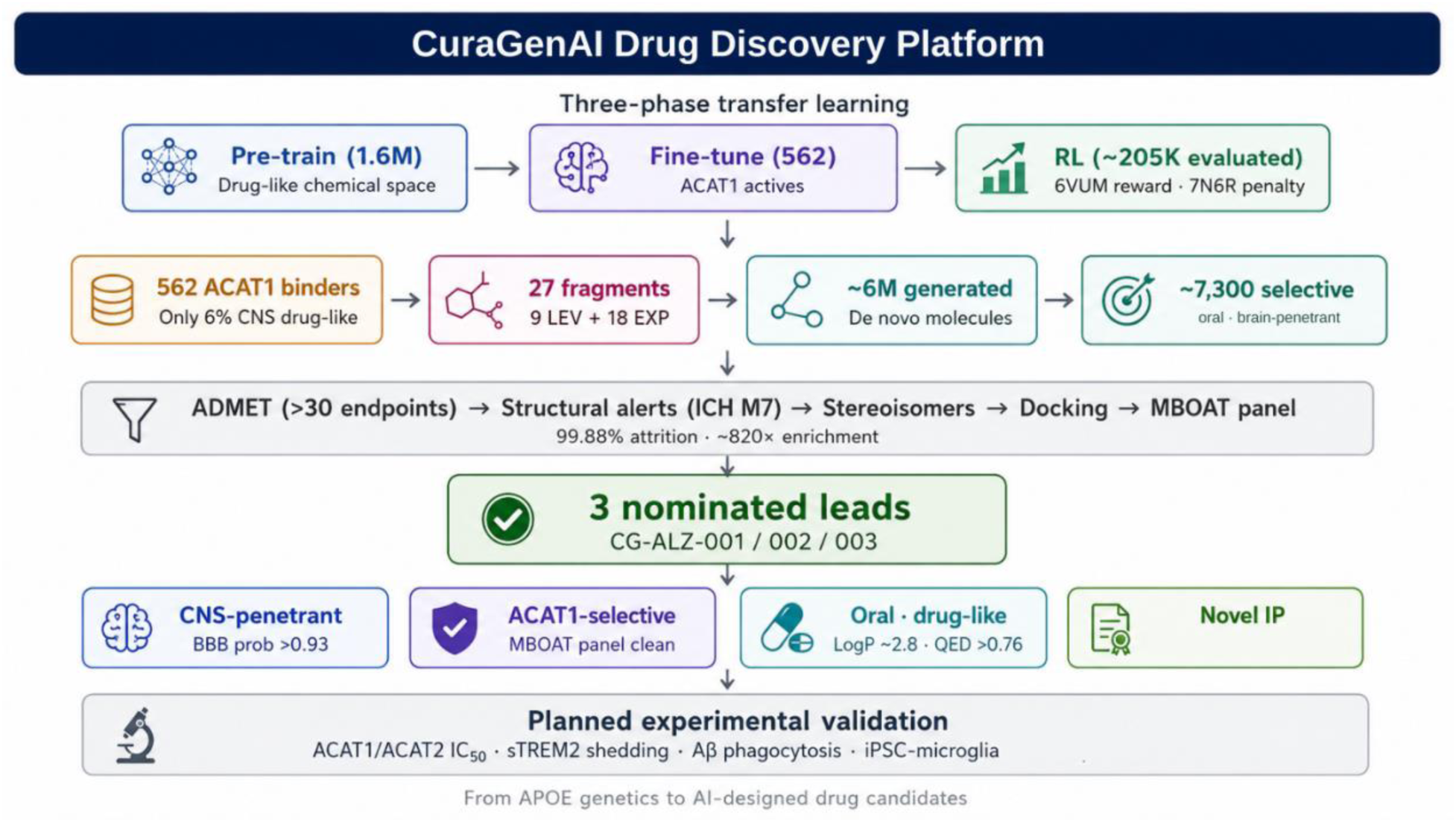
CuraGenAI Drug Discovery Platform overview. Three-phase transfer learning (pre-train on 1.6M drug-like molecules → fine-tune on 562 ACAT1 actives → RL with ACAT1 engagement and ACAT2 counter-screening over ∼205,000 evaluated molecules) drives generation of ∼6 million de novo candidates. Multi-stage filtering (ADMET, structural alerts, stereoisomers, docking, MBOAT counter-screening) yields ∼7,300 CNS-penetrant candidates (99.88% attrition) and three nominated leads. Planned experimental validation includes ACAT1/ACAT2 IC₅₀, iPSC-microglial proof-of-mechanism, and DMPK profiling.

**Figure 3.**
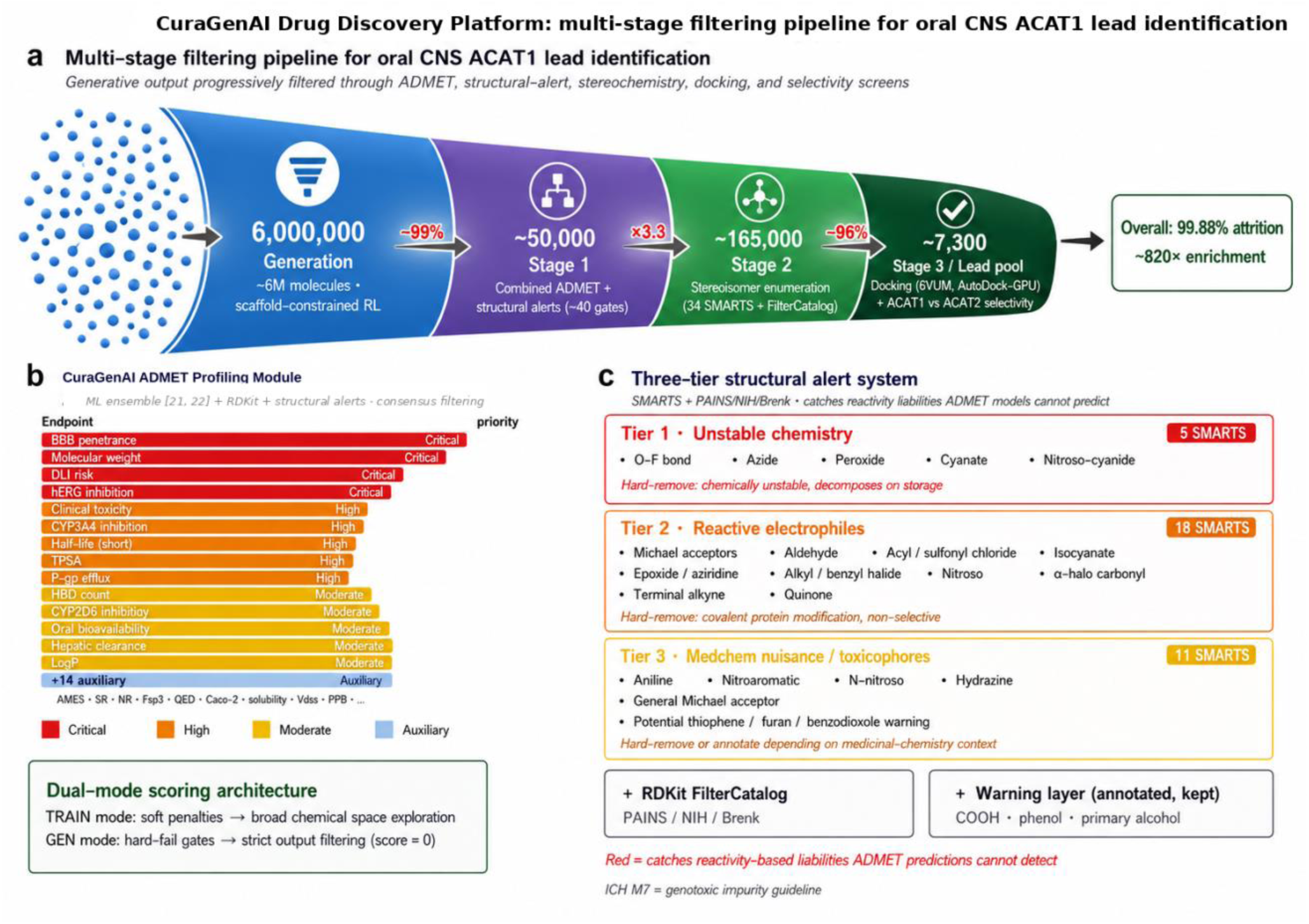
CuraGenAI Drug Discovery Platform: multi-stage filtering pipeline for oral CNS ACAT1 lead identification. (a) Generative output progressively filtered through physicochemical, ADMET, structural-alert, docking, and counter-screening steps, reducing approximately 6 million de novo-generated molecules to approximately 7,300 CNS-penetrant, ACAT1-targeted lead candidates. Overall attrition: 99.88%, enrichment: ∼820×. (b) CuraGenAI ADMET Profiling Module composite scoring with weighted endpoints and approximately 40 hard-fail gates. (c) Three-tier SMARTS-based structural alert system (5 unstable, 18 electrophilic, 11 medchem toxicophore patterns) complementing property-prediction models to catch reactivity-based liabilities.

Stage 1: Combined ADMET and structural alert hard-fail filtering applied a 28-endpoint weighted composite score with approximately 40 hard-fail gates, three-tier SMARTS-based structural alerts, and the RDKit FilterCatalog (PAINS, Brenk). CNS-optimized property gates included LogP ≤ 4.6, MW ≤ 451 Da, TPSA ≤ 80 Å², and HBD ≤ 2. The dual-mode scoring architecture: soft penalties during RL training, hard zero-scoring during generation, eliminated 99.1% of generated molecules at this stage alone.

Stage 2: Stereoisomer enumeration expanded ADMET-compliant parents into stereochemically defined entities (∼4× expansion). Ambiguous parents were dropped.

Stage 3: Structure-based docking and ACAT2 counter-screening (AutoDock-GPU) ranked candidates by predicted ACAT1 binding and enforced the ACAT1/ACAT2 selectivity gate.

Across the full pipeline, approximately 7,300 candidates passed ACAT1 engagement and ACAT2 counter-screening gates from approximately 6 million generated, an overall attrition rate of 99.88% (∼820-fold enrichment). Surviving candidate properties shifted significantly from the reference dataset: mean MW from 464 to approximately 385 Da, LogP from 5.3 to 2.8, TPSA from 81 to 65 Å², demonstrating that RL-guided generation combined with multi-stage filtering redirected chemical space from the lipophilic, peripherally-targeted profiles of historical ACAT1 inhibitors toward oral CNS drug-like space.

In effect, the integrated pipeline (combining scaffold-constrained generative chemistry, multi-objective RL optimization, and sequential ADMET/selectivity/safety filtering) performs the functional equivalent of iterative lead optimization, compressing what is traditionally a multi-year, resource-intensive medicinal chemistry campaign into a single computational workflow executed prior to synthesis. The three nominated leads described below enter experimental validation as pre-optimized development candidates rather than unrefined screening hits.

### 3.5 Predicted ACAT1 Binding of Lead Candidates

Lead candidates were evaluated by AutoDock Vina consensus docking (exhaustiveness 128, three seeds, median score) against two ACAT1 structures: PDB 6VUM (nevanimibe-bound) and PDB 6L47 (CI-976-bound). Docking scores were interpreted as relative binding energy estimates for compound ranking within the ACAT1 pocket, not as absolute binding affinity predictions. Nevanimibe (IC₅₀ = 9 nM ACAT1 vs 368 nM ACAT2, approximately 41-fold selectivity [41]) served as the ACAT1 reference benchmark, as the most advanced ACAT1-selective inhibitor with published enzymatic data.

CG-ALZ-001 and CG-ALZ-003 scored more favorably than nevanimibe (mean −9.16 and −9.26 vs −8.89 kcal/mol; Table 1), with consistent engagement across both ACAT1 structures. CG-ALZ-002 scored modestly weaker (−8.68, Δ = +0.21 kcal/mol vs nevanimibe), reflecting the multi-objective nature of the optimization, which balances target engagement against CNS penetrance, metabolic safety, and synthetic accessibility. All three candidates maintained stable binding poses in 50 ns molecular dynamics simulations (Section 3.7), supporting the docking-derived rankings.

**Table 1.** Predicted ACAT1 binding of CuraGenAI lead candidates vs nevanimibe (AutoDock Vina consensus docking; PDB 6VUM, 6L47). Scores in kcal/mol; more negative = stronger predicted binding.

| Compound | 6VUM | 6L47 | Mean ACAT1 | $\Delta$ vs Nevanimibe |
| --- | --- | --- | --- | --- |
| *Nevanimibe* | $-9.33$ | $-8.46$ | **$-8.89$** | — |
| **CG-ALZ-001** | $-9.51$ | $-8.81$ | **$-9.16$** | $-0.27$ |
| **CG-ALZ-002** | $-8.84$ | $-8.52$ | **$-8.68$** | $+0.21$ |
| **CG-ALZ-003** | $-9.58$ | $-8.94$ | **$-9.26$** | $-0.37$ |

### 3.6 MBOAT Family Counter-Screening

To assess predicted off-target liability across the MBOAT superfamily, all candidates passing ACAT1 docking and ADMET filters were counter-docked against four related enzymes: ACAT2 (PDB 7N6R, 7N6Q), DGAT1 (PDB 8ESM, 8ETM), PORCN (PDB 9OO6, 9OO7), and HHAT (PDB 7Q6Z, 7Q1U). Nevanimibe served as the selectivity anchor for all four targets based on its clinical safety at high oral exposures in Phase 1 and Phase 2 trials [42, 47] without signals attributable to MBOAT off-target inhibition. No lead candidate met the MBOAT counter-screening criterion for deprioritization. Because docking scoring functions are parameterized for within-pocket ligand ranking rather than cross-target affinity comparison, these assessments serve as a design-stage filter, not as quantitative selectivity predictions. Definitive isoform selectivity determination requires parallel IC₅₀ measurement, which is the first decision gate in the experimental validation campaign (Section 4.4).

### 3.7 Molecular Dynamics Validation of Binding Pose Stability

To confirm that docked poses represent stable binding modes, 50 ns explicit-solvent molecular dynamics simulations were performed for each lead candidate using OpenMM [45] with the AMBER ff19SB protein force field, GAFF2 ligand parametrization, TIP3P water, and 0.15 M NaCl. Following 5 ns of restrained equilibration, unrestrained production dynamics were analyzed for ligand heavy-atom RMSD and MM-GBSA binding free energy (Table 2).

**Table 2.** Binding pose stability from 50 ns molecular dynamics simulations.

| Compound | Ligand RMSD (Å) | MM-GBSA $\Delta G$ (kcal/mol) |
| --- | --- | --- |
| CG-ALZ-001 | 1.73 ± 0.16 | -28.91 ± 3.09 |
| CG-ALZ-002 | 0.72 ± 0.26 | -41.08 ± 5.77 |
| CG-ALZ-003 | 0.81 ± 0.15 | -34.58 ± 2.73 |

All three candidates maintained stable binding poses (RMSD < 2.5 Å) throughout the simulation, with favorable MM-GBSA binding free energies (−28.9 to −41.1 kcal/mol). CG-ALZ-002 and CG-ALZ-003 exhibited the lowest ligand RMSD values (0.72 and 0.81 Å respectively) and the most favorable predicted binding energies. Simulations were performed without an explicit lipid bilayer. Membrane-embedded simulations are planned as part of the experimental validation campaign.

### 3.8 Lead Candidate Profiles

Table 3 benchmarks the three leads against nevanimibe (selective ACAT1; discontinued due to adrenal toxicity) and avasimibe (non-selective ACAT1/ACAT2; withdrawn due to CYP3A4-mediated DDI and ACAT2-driven LDL elevation [18]). All three leads occupy CNS-compatible physicochemical space with zero Lipinski violations, contrasting sharply with both reference compounds (LogP >6.6, Lipinski failures). Predicted plasma protein binding of 78–84% (vs >99% for both references) suggests meaningfully improved free-fraction availability in the CNS compartment. Predicted half-lives of 6.5–12.9 h with moderate protein binding contrast with nevanimibe’s prolonged half-life (80.9 h), >99% PPB, and high lipophilicity, a profile conducive to adrenal tissue accumulation, consistent with its observed adrenal toxicity [41]. On safety endpoints, all leads fall below the 0.75 probability threshold across hERG, DILI, CYP3A4 inhibition, and carcinogenicity, whereas both references exceed this threshold on CYP3A4, consistent with the DDI liability that contributed to avasimibe’s withdrawal [18]. VEGA consensus mutagenicity classified all leads as non-mutagenic; nevanimibe returned a mutagenic classification. These profiles support advancement to synthesis and experimental IC₅₀ determination (Section 4.4).

**Table 3.** Predicted ADMET profiles of CuraGenAI lead candidates benchmarked against clinical-stage ACAT1 inhibitors. Physicochemical descriptors from RDKit; probability-based endpoints from ADMET-AI [21]; VEGA consensus mutagenicity from VEGA [43]. Green = within CNS target range; amber = borderline; red = outside range. Probability thresholds set at 0.75 per ADMET-AI model calibration.

| Property | CG-ALZ-001 | CG-ALZ-002 | CG-ALZ-003 | Nevanimibe | Avasimibe | Desirable range |
| --- | --- | --- | --- | --- | --- | --- |
| *Physicochemical* |  |  |  |  |  |  |
| MW (Da) | 365 | 389 | 390 | 422 | 502 | $\leq 450$ (CNS) |
| LogP (calc.) | 3.45 | 4.23 | 3.67 | 6.63 | 7.29 | $\leq 4.0$ (CNS) |
| TPSA ($\text{\AA}^2$) | 61.4 | 78.9 | 60.9 | 44.4 | 72.5 | $\leq 90$ (CNS) |
| HBD / HBA | 2 / 2 | 2 / 3 | 1 / 3 | 2 / 2 | 1 / 4 | $HBD \leq 3$ |
| Rotatable bonds | 4 | 6 | 5 | 7 | 10 | $\leq 8$ |
| QED | 0.87 | 0.76 | 0.85 | 0.53 | 0.37 | $> 0.5$ |
| Lipinski / Veber | Pass / Pass | Pass / Pass | Pass / Pass | Fail (LogP) / Pass | Fail ($\times 2$) / Pass | Pass / Pass |
| *CNS penetrance & oral PK* |  |  |  |  |  |  |
| BBB penetration, prob. | 0.96 | 0.93 | 0.98 | 0.95 | 0.38 | $> 0.85$ (CNS) |
| Bioavailability, prob. | 0.79 | 0.86 | 0.82 | 0.70 | 0.57 | $> 0.50$ |
| Solubility, log S | -3.4 | -3.4 | -3.8 | -6.2 | -5.2 | $> -5$ |
| PPB (% bound) | 79 | 78 | 84 | >99 | >99 | $< 95$ |
| Hepatocyte CL ($\mu\text{L}/\text{min}/10^6$) | 32.7 | 47.1 | 32.3 | 67.2 | 25.9 | Lower preferred |
| Half-life (h, Obach) | 9.4 | 12.9 | 6.5 | 80.9 | 111.9 | 8–24 h (QD) |
| *Safety & key liabilities* |  |  |  |  |  |  |
| hERG, prob. | 0.68 | 0.30 | 0.47 | 0.93 | 0.65 | $< 0.75$ |
| DILI, prob. | 0.23 | 0.24 | 0.47 | 0.36 | 0.60 | $< 0.75$ |
| CYP3A4 inhib., prob. | 0.30 | 0.61 | 0.10 | 0.82 | 0.82 | $< 0.75$ |
| Carcinogenicity, prob. | 0.08 | 0.09 | 0.07 | 0.41 | 0.82 | $< 0.75$ |
| VEGA Ames consensus | Non-mut. | Non-mut. | Non-mut. | Mutagenic | Non-mut. | Non-mut. |

## 4. Discussion

This work presents an AI-native pipeline for the de novo design of oral, brain-penetrant ACAT1 inhibitor candidates for Alzheimer’s disease. Multi-stage filtering of approximately 6 million generated molecules yielded three lead candidates with favorable predicted CNS, counter-screening, and safety profiles. The following sections assess their significance, differentiation, and translational path.

### 4.1 Significance and Therapeutic Context

This work translates foundational ACAT1/SOAT1 biology [9, 10], preclinical AD studies linking ACAT1 inhibition to amyloid and microglial mechanisms [11, 12], and recent ACAT1 structural biology [25] into a CNS-focused small-molecule design strategy. The convergence of human genetics [4], microglial lipid biology [7, 12], and AI-enabled drug design positions ACAT1 as an attractive therapeutic target for Alzheimer’s disease. The present work demonstrates that structure-based reinforcement learning on experimentally determined cryo-EM structures can generate drug-like ACAT1-targeted candidates with predicted properties compatible with chronic oral CNS dosing. Importantly, the resulting leads occupy physicochemical space distinct from historical ACAT inhibitor series, with lower lipophilicity and improved predicted CNS drug-like profiles.

### 4.2 Comparison to Alternative Computational Approaches

The CuraGenAI pipeline differs from conventional AI-driven drug discovery in its reliance on structure-based docking as the primary RL reward signal rather than QSAR surrogate models. Most published generative platforms (including GENTRL, REINVENT, and related approaches) train QSAR predictors on large bioactivity datasets (typically >5,000 compounds) and score generated molecules against these surrogates. This is effective for well-characterized targets with abundant data (kinases, GPCRs, proteases) but presents a practical limitation for underexplored targets where small training sets produce QSAR artifacts that the RL agent optimizes against.

ACAT1 exemplifies this challenge: public bioactivity data are insufficient for training a robust QSAR predictor, particularly one distinguishing ACAT1 from ACAT2. Our approach handles this by using the curated 562-compound set exclusively for fine-tuning the generative model, while the RL reward signal is derived from physics-based docking (AutoDock-GPU [46] against PDB 6VUM). This structure-based reward directly evaluates each generated molecule’s complementarity to the ACAT1 catalytic pocket rather than relying on a statistical approximation learned from limited training data. Although physics-based docking is inherently slower than QSAR inference, GPU-accelerated docking combined with parallelized workflow design maintained practical throughput without requiring specialized HPC infrastructure.

An alternative approach, high-throughput virtual screening of large commercial libraries such as ZINC (>1 billion compounds) or Enamine REAL (>6 billion), was considered but not selected for two reasons. First, virtual screening ranks existing compounds by predicted binding but does not generate novel chemical matter, limiting the ability to explore regions of chemical space not represented in commercial catalogs. Second, virtual screening provides no feedback between scoring objectives: molecules are ranked by binding score alone, and ADMET filtering is applied post hoc to a naive population, typically eliminating over 95% of hits. By contrast, the CuraGenAI RL reward function integrates binding, ADMET, and selectivity signals during training, so each generation cycle produces molecules progressively enriched for multi-objective compliance before post-generation filtering is applied. The pipeline is further differentiated by dual-source fragment curation seeding the generative model with target-relevant, CNS-compatible chemical matter (Section 2.5) and, to our knowledge, the first structured MBOAT family counter-screening panel for ACAT1 inhibitor design, a practice we recommend as standard for any ACAT1 program.

### 4.3 Overcoming Historical Failure Modes

Every prior ACAT1 program was discontinued before demonstrating clinical efficacy, but each for specific, identifiable reasons that now inform our design. Understanding these failure modes and how the current pipeline addresses them is central to assessing candidate viability.

#### 4.3.1 Clinical Evidence for ACAT1 Target Safety

The clinical evidence for ACAT1 inhibitor safety is more reassuring than the preclinical literature suggests. In the Phase 1 adrenocortical carcinoma trial, 63 patients received oral nevanimibe at doses up to approximately 11,000 mg/day, yet only two (3.2%) experienced adrenal insufficiency, classified as on-target pharmacology in diseased adrenal tissue rather than off-target toxicity [42]. In the Phase 2 congenital adrenal hyperplasia study (125–1,000 mg twice daily), no dose-related adrenal insufficiency was observed and a maximum tolerated dose could not be defined [47], indicating a wide tolerability window at clinical exposures. The adrenal liability is an on-target, dose-dependent phenomenon driven by cholesteryl ester depletion in adrenal tissue, compounded by nevanimibe’s pharmacokinetic persistence (predicted half-life ∼81 h). Critically, ACAT1 expression is significantly elevated in AD microglia [12, 13], with cholesteryl ester levels increased approximately 2-fold in AD brain tissue [13]. The therapeutic objective is therefore partial normalization of pathologically upregulated ACAT1 activity, a fundamentally lower-exposure strategy than the near-total suppression targeted in oncology.

#### 4.3.2 CuraGenAI Design Response to Historical Failure Modes

Our pipeline targets each historical failure mode directly. CNS-optimized physicochemical filters produce leads with substantially lower lipophilicity than prior compounds (LogP ∼2.8 vs 5.3), reducing the peripheral tissue accumulation underlying nevanimibe’s adrenal toxicity. The selectivity-by-design strategy, incorporating an ACAT2 inverse docking penalty during RL training and broader MBOAT off-target filtering, is designed to avoid the hepatic cholesteryl ester depletion and LDL elevation that led to the discontinuation of the avasimibe and pactimibe programs. CYP3A4 inhibition, the specific DDI liability that contributed to avasimibe’s withdrawal, is penalized during both RL training and hard-fail filtering, and all three leads score below the 0.75 probability threshold. Finally, BBB penetrance optimization ensures predicted CNS exposure (BBB probability >0.93 for all leads), directly addressing a gap in brain-targeted design across prior ACAT1 programs.

### 4.4 Clinical Positioning and Translational Path

An oral, brain-penetrant, ACAT1-targeted small molecule would address Aβ clearance, Aβ production, and tau degradation through a single mechanism without the ARIA risk associated with anti-amyloid antibodies. This offers a differentiated and potentially complementary approach to current therapies, which target deposited plaque but do not engage the upstream cholesterol-driven pathology explored here. Experimental validation will proceed through a staged, decision-gated sequence: synthesis and enzymatic ACAT1 IC₅₀, isoform selectivity confirmation, induced pluripotent stem cell (iPSC)-microglial proof-of-mechanism [12], drug metabolism and pharmacokinetics (DMPK) profiling with brain exposure measurement, and new approach methodologies (NAM) safety confirmation (hepatic spheroids, hERG, AI-augmented toxicity predictions), with the goal of assembling an IND-enabling data package. Also within the platform, safety and selectivity constraints are not applied solely as post-generation filters but are integrated directly into the RL reward function, so each generation cycle produces molecules pre-optimized for these endpoints. This NAM-forward strategy aligns with the FDA Modernization Act 3.0 [48], the FDA’s Roadmap to Reducing Animal Testing [49], and the March 2026 FDA draft guidance on NAM validation [50].

### 4.5 Limitations

Docking scores are computational estimates for compound ranking, not absolute binding affinity predictions. MBOAT counter-screening analysis uses rigid-body docking rather than free energy perturbation (FEP) or molecular dynamics (MD). Machine learning ADMET predictions have applicability domain limitations, especially for novel chemotypes not well-represented in training data, and the orthogonal safety confirmation strategy cannot fully substitute for experimental testing. All computational predictions require experimental confirmation through the staged validation program described in Section 4.4.

## 5. Conclusion

Alzheimer’s disease remains among the most pressing unmet needs in neurology. The convergence of human genetic evidence spanning common APOE allelic variation (estimated to account for 72–93% of AD burden across cohorts [4]), TREM2 risk variants, and cholesterol transport polymorphisms, implicates microglial lipid handling as a central driver of disease pathology. ACAT1 occupies an attractive position within this axis: its inhibition simultaneously increases sTREM2-dependent Aβ phagocytosis [12], reduces Aβ production [11], and promotes phospho-tau degradation via autophagy [14], targeting amyloid, tau, and neuroinflammation through a single target without the ARIA risk of anti-amyloid antibodies.

We present the CuraGenAI pipeline for the design of oral, brain-penetrant ACAT1 inhibitors, integrating structure-based reinforcement learning with selectivity-by-design, multi-objective ADMET optimization, and MBOAT family counter-screening. From approximately 6 million de novo-generated molecules, multi-stage filtering achieved 99.88% attrition, yielding approximately 7,300 CNS-penetrant, ACAT1-targeted candidates with favorable predicted oral CNS profiles (MW 350–420 Da, LogP 2.0–3.5, TPSA 50–75 Å²) and synthetic accessibility suitable for synthesis. Of these, three lead candidates have been prioritized for synthesis, parallel ACAT1/ACAT2 IC₅₀ determination, and pharmacokinetic profiling, with the goal of establishing experimentally validated, ACAT1-selective, brain-penetrant candidates for Alzheimer’s disease.

Beyond this program, recent advances in generative AI, GPU-accelerated molecular simulation, and machine learning safety prediction have compressed what traditionally required years and tens of millions of dollars into workflows that are faster, more cost-effective, and designed to reduce late-stage attrition. If programs like this succeed in translating computational predictions into experimentally validated clinical candidates, they may expand therapeutic options available for more than 55 million people worldwide living with dementia [1], for whom effective disease-modifying treatment remains an urgent, unmet need.

## Author Contributions

S.A. conceived the project, designed the platform architecture and computational pipeline, performed all computational experiments, analyzed the data, and wrote the manuscript. R.P. contributed to strategic platform positioning, manuscript figure design, and critical review of platform architecture. P.A. provided critical review of the manuscript and computational methodology. C.R. provided critical review of the biological rationale and translational interpretation. S.G. contributed clinical and translational review of the Alzheimer’s disease rationale and manuscript.

## Acknowledgments

The authors thank Dr. Ta-Yuan Chang and Dr. Catherine C.Y. Chang (Dartmouth College) for their foundational work on ACAT1 biochemistry and selectivity that informed the fragment curation and chemotype classification in this study. We acknowledge the structural biology contributions of Dr. Tobias Long and colleagues (UT Southwestern) whose cryo-EM structure of nevanimibe-bound ACAT1 (PDB 6VUM) and Dr. Chengcheng Guan, Dr. Lei Chen and colleagues (Peking University) whose CI-976-bound ACAT1 structure (PDB 6L47) enabled the structure-based design program. We thank Prof. Rudolph E. Tanzi and Moriah J. Hovde (Massachusetts General Hospital) for their discovery of the ACAT1-sTREM2-LRP1 pathway that provided the mechanistic foundation for this work. We acknowledge Dr. Gerard J.P. van Westen and colleagues (Leiden University) for the open-source generative chemistry tools that informed the development of the CuraGenAI pipeline. We thank Parker Moss for strategic guidance on platform positioning.

## Data Availability

Fragment classification, scaffold classes, and physicochemical properties are provided in Supplementary Table S2. Fragment structures and the 562-compound dataset are available from the corresponding author upon reasonable request.

## Supplementary Methods: Computational Protocol Details

This section provides complete software versions, docking parameters, receptor preparation protocols, and box coordinates to enable independent reproduction of the structure-based scoring described in the main text.

### S1. Software Versions

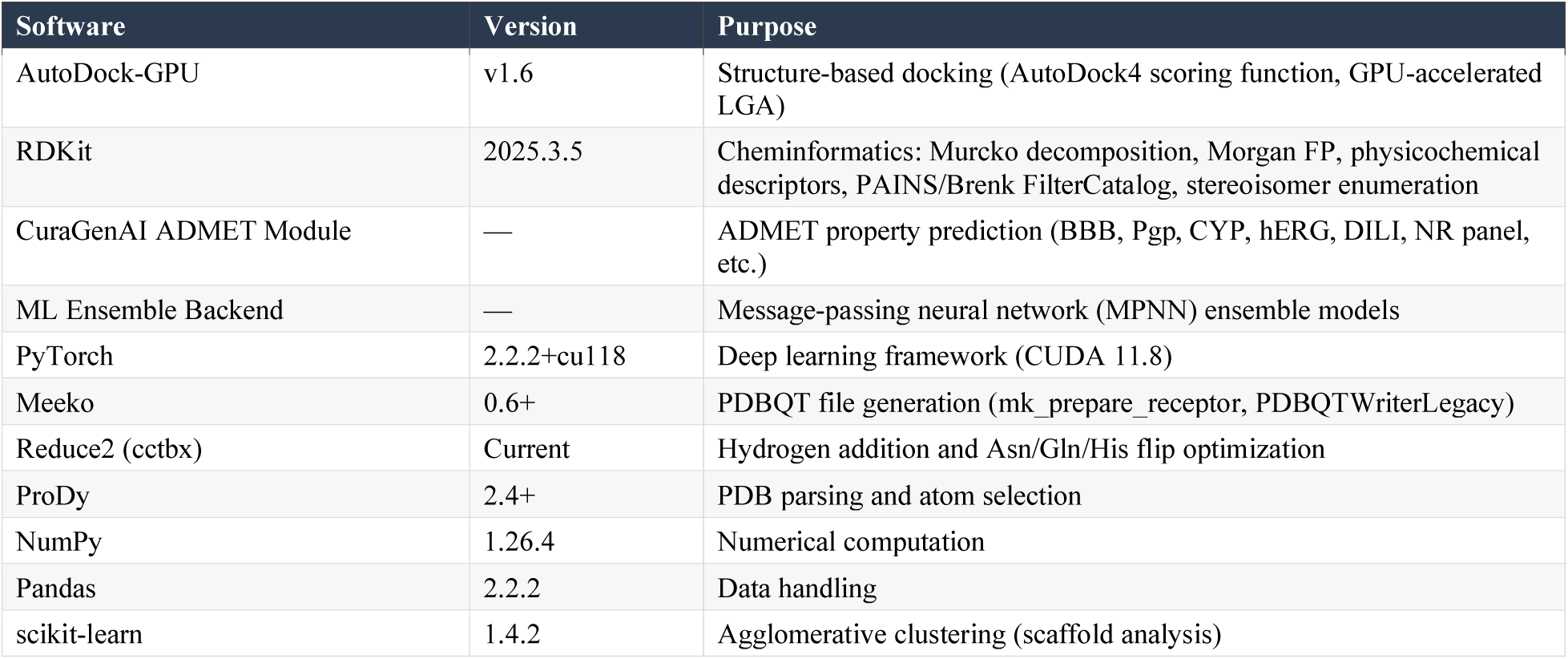

| Software | Version | Purpose |
| --- | --- | --- |
| AutoDock-GPU | v1.6 | Structure-based docking (AutoDock4 scoring function, GPU-accelerated LGA) |
| RDKit | 2025.3.5 | Cheminformatics: Murcko decomposition, Morgan FP, physicochemical descriptors, PAINS/Brenk FilterCatalog, stereoisomer enumeration |
| CuraGenAI ADMET Module | — | ADMET property prediction (BBB, Pgp, CYP, hERG, DILI, NR panel, etc.) |
| ML Ensemble Backend | — | Message-passing neural network (MPNN) ensemble models |
| PyTorch | 2.2.2+cu118 | Deep learning framework (CUDA 11.8) |
| Meeko | 0.6+ | PDBQT file generation (mk_prepare_receptor, PDBQTWriterLegacy) |
| Reduce2 (cctbx) | Current | Hydrogen addition and Asn/Gln/His flip optimization |
| ProDy | 2.4+ | PDB parsing and atom selection |
| NumPy | 1.26.4 | Numerical computation |
| Pandas | 2.2.2 | Data handling |
| scikit-learn | 1.4.2 | Agglomerative clustering (scaffold analysis) |

### S2. AutoDock-GPU Docking Parameters

| Parameter | Value | Rationale |
| --- | --- | --- |
| Number of LGA runs | `-n 64` | 64 independent runs per ligand; substantially more exhaustive than default (10–20) to ensure convergence in the deep TM4–TM9 transmembrane pocket |
| Energy evaluations per run | `--nev 20000000` | 20 million evaluations; 8× the default (2.5M). Required for adequate sampling of large, flexible ligands (MW 350–450, 5–10 rotatable bonds) |
| Autostop | `--autostop 0` | Disabled. Early stopping can prematurely terminate runs in buried membrane pockets where initial convergence is slower |
| Random seed | `--seed 42` | Fixed seed for reproducibility across generation blocks |
| Scoring function | AutoDock4 | Physics-based scoring (van der Waals, H-bonding, electrostatic, desolvation, torsional penalty) |
| GPU work items | 64wi (128wi build) | NVIDIA L4 Tensor Core GPU (24 GB VRAM, CUDA 12) |
| Pose selection | Global best | Lowest binding energy across all runs and all protonation/tautomeric states evaluated per molecule |

### S2B. AutoDock Vina Lead Rescoring Parameters

Final lead candidates were rescored using AutoDock Vina with exhaustiveness 128 across three independent random seeds. The reported score is the median best-pose score across seeds. Scores were interpreted as relative ranking metrics within the same receptor pocket.

### S3. Hardware and Throughput

All docking and molecular generation computations were performed on NVIDIA L4 Tensor Core GPUs (24 GB VRAM) via Google Cloud Platform with CUDA 12.

**Supplementary Table S1:** ADMET Scoring Endpoint Categories and Clinical Rationale. The following table provides the endpoint categories used in the CuraGenAI ADMET Profiling Module, with threshold categorization and clinical rationale. Scoring parameters were derived from published CNS drug space boundaries [34] and ACAT1-specific safety considerations [18].

| Endpoint | Transfer Fn | Threshold Cat. | Weight Cat. | Clinical Rationale / Basis |
| --- | --- | --- | --- | --- |
| BBB permeability | Direct prob. | Stringent | High | No prior program achieved CNS exposure |
| P-gp efflux | $1 - p$ | Stringent | High | #1 barrier to CNS drug delivery |
| TPSA | Gaussian | Stringent | High | CNS MPO [34]; strong BBB correlate |
| HBD count | Gaussian | Moderate | Moderate | Each HBD reduces BBB ~10-fold |
| Molecular weight | Gaussian | Stringent | High | CNS drugs typically <450 Da |
| LogP | Gaussian | Moderate | Moderate | CNS range 1–4; >4 causes accumulation |
| hERG inhibition | Sigmoid inv. | Stringent | High | ICH E14; chronic dosing requires clean hERG |
| DILI risk | 1 – p | Stringent | High | ACAT1 hepatic expression; chronic use |
| Clinical toxicity | 1 – p | Stringent | High | Composite clinical toxicity predictor |
| Carcinogenicity | 1 – p | Moderate | Moderate | Chronic 5–10+ year AD therapy |
| Ames mutagenicity | Hard fail only | Stringent | Gen-mode only | ICH S2; non-negotiable for chronic therapy |
| CYP3A4 inhibition | 1 – p | Stringent | High | Avasimibe CYP3A4 induction caused failure |
| CYP2D6 inhibition | 1 – p | Moderate | Moderate | AD patients on donepezil (CYP2D6 substrate) |
| CYP2C9 inhibition | 1 – p | Moderate | Moderate | Polypharmacy coverage |
| CYP2C19 inhibition | 1 – p | Moderate | Moderate | Polypharmacy coverage |
| CYP1A2 inhibition | 1 – p | Moderate | Moderate | Polypharmacy coverage |
| CYP3A4_Substrate | Penalty | Moderate | Penalty | Metabolic DDI vulnerability |
| Half-life | Gaussian | Moderate | High | Short t<sub>1/2</sub> for systemic-sparing |
| Clearance_Hepatocyte | Gaussian | Moderate | Moderate | Moderate-high CL avoids accumulation |
| Volume of distribution | Gaussian | Moderate | Moderate | Low-moderate V<sub>d</sub> for adrenal sparing |
| Plasma protein binding | Gaussian | Moderate | Moderate | >95% binding reduces free brain fraction |
| Oral bioavailability | Direct prob. | Permissive | Moderate | Adequate oral absorption |
| Intestinal absorption | Direct prob. | Permissive | Moderate | Intestinal absorption proxy |
| Caco2 Wang | Sigmoid | Permissive | Moderate | Permeability proxy |
| Aqueous solubility | Sigmoid | Moderate | Moderate | Oral absorption requirement |
| Fsp3 | Gaussian | Moderate | Moderate | 3D character; clinical success [33] |
| QED | Direct score | Permissive | Low | Composite drug-likeness [29] |
| Rotatable bonds | Gaussian | Moderate | Low | CNS drugs typically <7 |
| NR: Androgen receptor | 1 – p | Moderate | Moderate | Endocrine safety; chronic dosing |
| NR: Estrogen receptor | 1 – p | Moderate | Default | Endocrine safety |
| NR: Aromatase | 1 – p | Moderate | Default | Endocrine safety |
| NR: PPAR-γ | 1 – p | Moderate | Default | Endocrine safety |
| NR-AhR | 1 – p | Moderate | Penalty | CYP induction / DDI proxy |
| SR: MMP | Hard fail | Moderate | Default | Mitochondrial toxicity marker |
| SR: p53 | Hard fail | Moderate | Default | Genotoxicity marker |
| SR: HSE | Hard fail | Moderate | Default | Heat shock / protein stress |
| SR: ARE | Hard fail | Moderate | Default | Oxidative stress marker |
| SR: ATAD5 | Hard fail | Moderate | Default | DNA damage marker |
| PAINS/Brenk alerts | Binary | Stringent | Penalty/Fail | Reactive/promiscuous compounds [30, 31] |

Transfer function types: Gaussian = centered desirability; Sigmoid = logistic transformation; Safety inversion = higher liability → lower score; Direct = used as-is; Hard fail = binary elimination in generation mode. Threshold categories: Stringent = critical for CNS/safety; Moderate = important but with tolerance; Permissive = supplementary quality metric. Weight categories: High, Moderate, Low, and Default reflect endpoint clinical relevance; Penalty = applied as multiplicative penalty; Gen-mode only = hard fail in generation mode, not scored during training.

**Supplementary Table S2.** Fragment library: classification, physicochemical properties, and design rationale (27 fragments). Source A = data-driven Murcko extraction; Source B = expert curation. LEVERAGE = validated ACAT1 pharmacophores; EXPLORE = novel CNS-privileged scaffolds.

| # | Name | Class | Source | Scaffold | MW | LogP | TPSA | Design Rationale |
| --- | --- | --- | --- | --- | --- | --- | --- | --- |
| 1 | DataDriven-Novel-1 | EXPLORE | Data | Arene | 78 | 1.7 | 0.0 | Minimal aromatic seed from top-scoring novel family |
| 2 | DataDriven-Novel-2 | EXPLORE | Data | Diarylethane | 182 | 3.5 | 0.0 | Hydrophobic linker from top-scoring novel family |
| 3 | DataDriven-Novel-3 | EXPLORE | Data | Phenylcyclopentyl amide | 294 | 4.3 | 41.1 | Branched amide scaffold; 5 dataset members |
| 4 | DataDriven-F12511 | LEVERAGE | Data | Dihydrazide | 268 | 1.6 | 58.2 | Top F12511 family scaffold |
| 5 | DataDriven-Piperidine | LEVERAGE | Data | Piperidine-pyridine urea | 296 | 2.9 | 57.3 | Top piperidine family; CNS linker motif |
| 6 | DataDriven-Chromanone | EXPLORE | Data | Chromone-chromene | 278 | 3.8 | 35.5 | Top chromanone/flavanone family scaffold |
| 7 | K-604 core | LEVERAGE | Expert | Benzimidazole | 175 | 1.5 | 57.8 | 229× ACAT1 selectivity. NH→Asn421. Patent expired 2019 |
| 8 | K-604 bidirectional | LEVERAGE | Expert | Benzimidazole | 175 | 0.6 | 71.8 | Two exit vectors: His460 + cholesterol tunnel |
| 9 | K-604 tail | LEVERAGE | Expert | Methylthiopyridine | 242 | 2.8 | 42.0 | Bis-SCH<sub>3</sub> engages TM7/TM8; key selectivity determinant |
| 10 | CI-976 headgroup | LEVERAGE | Expert | Trimethoxyphenyl | 225 | 1.7 | 56.8 | Cryo-EM co-crystal (6L47). Validated binding mode |
| 11 | F12511 headgroup | LEVERAGE | Expert | Di-tert-butylphenol | 263 | 3.0 | 63.3 | Most potent ACAT1 inhibitor (IC<sub>50</sub> = 39nM). Patent expired |
| 12 | CF<sub>3</sub>-pyridine | LEVERAGE | Expert | Trifluoromethylpyridine | 204 | 2.1 | 42.0 | Novel CF<sub>3</sub> bioisostere of K-604 SCH<sub>3</sub>. No existing patents |
| 13 | 4-Phenylpiperidine | LEVERAGE | Expert | Phenylpiperidine | 161 | 2.2 | 12.0 | Classical CNS linker (donepezil). Proven BBB uptake |
| 14 | Benzothiazole amide | EXPLORE | Expert | Benzothiazole | 192 | 1.3 | 56.0 | NH→S swap from K-604. Reduces HBD by 1. CNS scaffold |
| 15 | Indole acetamide | EXPLORE | Expert | Indole | 174 | 2.1 | 44.9 | CNS-privileged. Indole NH mimics benzimidazole→Asn421 |
| 16 | Chromanone | EXPLORE | Expert | Chromanone | 148 | 1.2 | 26.3 | Glabrol-derived (natural ACAT1 inhibitor). Low toxicity |
| 17 | Quinolinone | EXPLORE | Expert | Quinolinone | 145 | 1.5 | 32.9 | C=O + NH engage His460/Asn421. Brain-penetrant |
| 18 | Coumarin acetamide | EXPLORE | Expert | Coumarin | 203 | 1.8 | 59.3 | Mimics cholesterol ring A/B in lateral tunnel (6VUM) |
| 19 | Isoflavan core | EXPLORE | Expert | Isoflavan | 210 | 3.8 | 9.2 | Glabridin-derived. Chiral C3 for pocket asymmetry |
| 20 | Oxindole | EXPLORE | Expert | Oxindole | 133 | 1.2 | 29.1 | NP-like lactam. C3 sp<sup>3</sup> adds 3D. CNS (sunitinib) |
| 21 | Benzofuran amide | EXPLORE | Expert | Benzofuran | 161 | 1.5 | 56.2 | Neutral H-bonding reduces P-gp efflux risk |
| 22 | Dihydrocoumarin | EXPLORE | Expert | Dihydrocoumarin | 148 | 1.4 | 26.3 | Esterase-labile lactone for shorter systemic exposure |
| 23 | Stilbene | EXPLORE | Expert | Stilbene | 240 | 3.9 | 18.5 | Pterostilbene-type. Superior BBB vs resveratrol |
| 24 | Xanthone | EXPLORE | Expert | Xanthone | 196 | 2.9 | 30.2 | Membrane-interactive NP. Engages TM4–TM9 opening |
| 25 | N-Phenyl carbamate | EXPLORE | Expert | Carbamate | 152 | 0.7 | 78.3 | Soft-drug. Esterase-labile prevents accumulation |
| 26 | Cyclopentyl urea | EXPLORE | Expert | Urea | 232 | 3.4 | 41.1 | Nevanimibe minimal pharmacophore without acyl chains |
| 27 | Homovanillyl acetamide | EXPLORE | Expert | Vanilloid | 195 | 1.0 | 58.6 | Capsaicin-inspired. Phenol+methoxy for ER membrane |

**Summary statistics.** 27 fragments total: 9 LEVERAGE + 18 EXPLORE. Source: 6 data-driven + 21 expert-curated. Mean MW = 199 Da (range 78–296), mean LogP = 2.2 (range 0.6–4.3), mean TPSA = 40.9 Å<sup>2</sup> (range 0.0–78.3). For comparison, the 562-compound reference dataset has mean MW = 464 Da, mean LogP = 5.3, mean TPSA = 81.0 Å<sup>2</sup>.
**Abbreviations:** BBB = blood–brain barrier; CNS = central nervous system; CoM = composition-of-matter; HBD = hydrogen bond donor; NP = natural product; SAR = structure-activity relationship.

**Supplementary Table S3.** Docking box coordinates for ACAT1 and ACAT2 structures. All boxes defined by enveloping the co-crystallized inhibitor with 5 Å padding using Meeko mk_prepare_receptor. Coordinates in the PDB crystal frame.

| Target | PDB | Center X | Center Y | Center Z | Size X (Å) | Size Y (Å) | Size Z (Å) | Ref. Ligand |
| --- | --- | --- | --- | --- | --- | --- | --- | --- |
| ACAT1 (primary) | 6VUM | 158.234 | 152.897 | 145.321 | 19.138 | 15.845 | 21.981 | Nevanimibe (ROV) |
| ACAT1 (validation) | 6L47 | 85.609 | 98.865 | 104.071 | 18.914 | 18.660 | 20.527 | CI-976 (E5L) |
| ACAT2 (primary) | 7N6R | 149.507 | 113.636 | 123.759 | 18.559 | 18.224 | 20.002 | Nevanimibe (ROV) |
| ACAT2 (validation) | 7N6Q | 137.726 | 110.119 | 125.518 | 21.180 | 19.121 | 21.984 | PPPA (7T8) |

Docking box coordinates and scores for MBOAT off-targets (DGAT1, PORCN, HHAT) are available from the corresponding author upon reasonable request.

